# PLANET-MD: Ultra-fast Proteome-scale Prediction of Allosteric Networks in Proteins

**DOI:** 10.1101/2025.06.12.659353

**Authors:** Samuel Sledzieski, Sonya M. Hanson

## Abstract

Proteins are dynamic molecules that depend on conformational flexibility to carry out functions in the cell, yet despite significant advances in the modeling of static protein structure, prediction of these dynamics remains challenging. We introduce PLANET-MD, a machine learning model that predicts dynamic protein properties from sequence or static structure with unprecedented speed and accuracy. Trained on thousands of molecular dynamics trajectories spanning diverse protein families, PLANET-MD simultaneously models multiple dynamics features: root-mean-square fluctuations (RMSF), generalized correlation coefficients (GCC-LMI), and a novel structural heterogeneity profile (SHP) based on recent structure quantization methods. PLANET-MD significantly outperforms existing methods in predicting simulation-derived dynamics. We reduce RMSF prediction error by 57% compared to BioEmu and calibrated Dyna-1 predictions, including an up to 73% error reduction for long proteins. We validate these predictions with experimental hetNOE data, and we demonstrate the ability to adapt predictions to different physical temperatures. We highlight PLANET-MD’s utility in constructing allosteric networks in the oncogene *KRAS* and identify structural sub-modules with correlated motions, and we validate PLANET-MD by showing that changes in node centrality within predicted *KRAS* allosteric networks correlate with changes of folding free energy in experimental DMS data. Our approach makes predictions in seconds rather than hours or days, enabling us to perform the first comprehensive dynamics analysis of the entire human proteome. PLANET-MD bridges the gap between static structural biology and dynamic functional understanding, enabling dynamics-aware structural analysis and variant effect prediction at scales previously unavailable. PLANET-MD is available as free and open-source software at https://github.com/flatironinstitute/PLANET-MD.

---

Proteins are not static entities but dynamic molecules that adopt diverse conformational states essential to their biological function. This conformational flexibility is critical for processes ranging from enzyme catalysis and allosteric regulation to protein-protein interactions and molecular recognition. While machine learning for structural biology has made remarkable progress in determining high-resolution protein structures (Jumper et al., 2021; Wu et al., 2022; Lin et al., 2023), these represent snapshots of ensembles and miss the dynamics that determine protein function. To build a better understanding of how a protein functions, it is crucial to consider the motion of its structure. Further, it has been established that allostery is a widespread property of dynamic proteins (Gunasekaran, Ma, and Nussinov, 2004; Lindsley and Rutter, 2006; Astore et al., 2024; Liao and Lehner, 2025), and recent studies have begun to model not just dynamics, but the mechanism by which allostery propagates through proteins (Bowerman and Wereszczynski, 2016; Ahuja, Taylor, and Kornev, 2019; Weng et al., 2024; Beltran et al., 2026).

While molecular dynamics (MD) simulations that provide atomic-resolution trajectories have traditionally been the gold standard for exploring protein dynamics, such simulations are computationally expensive and typically require days to weeks of dedicated computing to generate a single trajectory of sufficient length. This computational limitation prevents systematic proteome-scale analysis of allostery and limits the exploration of sequence variants, hampering applications in protein engineering, drug discovery, and understanding of disease mechanisms. Still, recent work in machine learning for proteins points to an opportunity for scalable modeling of the structural dynamics that underlie protein function. Large protein sequence data sets (Suzek et al., 2015; Steinegger and Söding, 2018) enabled the training of protein language models (Rives et al., 2019; Bepler and Berger, 2021), and these models combined with large protein structure data sets (Burley et al., 2023) enabled highly accurate protein structure models (Wu et al., 2022; Lin et al., 2023). The recent introduction of large scale molecular dynamics data sets (Schweke et al., 2023; Mirarchi, Giorgino, and De Fabritiis, 2024) could provide a similar opportunity to scale computational modeling of protein dynamics and allostery using machine learning (Sledzieski and Hanson, 2026).

Here, we introduce PLANET-MD (**P**redicting **L**ocal fluctuations and **A**llosteric **N**etworks via **E**mbedding **T**rajectories from **M**olecular **D**ynamics), a machine learning approach that combines protein sequence and structure pretraining with supervision on molecular dynamics trajectories to model allosteric behavior at the scale of the proteome. A key motivation of this work is prior research showing that nanosecond-scale equilibrium fluctuations of a protein may be sufficient to infer allosteric networks and communication (Kern and Zuiderweg, 2003; Henzler-Wildman et al., 2007; Hammes, Chang, and Oas, 2009; Singh and Bowman, 2017; Maschietto et al., 2023). We thus propose that short-duration MD simulations that capture such fluctuations contain sufficient information to train a model to predict pairwise correlations and allosteric networks—even as these simulations may be biased by initial configuration and fail to access larger conformational changes. Two recently-released MD data sets, ATLAS (Schweke et al., 2023) and mdCATH (Mirarchi, Giorgino, and De Fabritiis, 2024) provide exactly this substrate, and motivate our work here (Figure 1A). Trained on thousands of simulation trajectories from diverse sequences and structural folds, PLANET-MD leverages recent advances in protein language models and structure quantization to encode these trajectories into compressed information-dense representations that are predictable from sequence (Methods, Figure 1B). We predict three distinct quantities: root-mean-square fluctuations (RMSF), generalized correlation coefficients with linear mutual information (GCC-LMI), and a newly-developed structural heterogeneity profile (SHP), all of which enable characterization of functional dynamics and inference of allosteric networks (Figure 1C).

PLANET-MD offers several key advantages: (1) accurate recovery of protein dynamic properties; (2) flexibility in input requirements, functioning with sequence alone or optionally augmented with a structure; and (3) ultra-fast prediction, reducing computation time from hours or days to fractions of a second and enabling the first comprehensive survey of protein dynamics across the entire human proteome (Figure 7). These capabilities open new possibilities for identifying proteins with similar dynamic properties, discovering cryptic binding sites, modeling allosteric communication networks, and predicting the functional impact of sequence variants. We anticipate that PLANET-MD will be complementary to existing sampling-based methods (Lewis et al., 2025; Liu et al., 2025), enabling capabilities such as guided sampling in regions of high uncertainty. Our work bridges the gap between static structural biology and a dynamic model of protein function, providing a computational tool that complements experimental and sampling approaches and enhancing our ability to explore the relationship between protein sequence, structure, and allostery at unprecedented scales.

**Figure 1:**
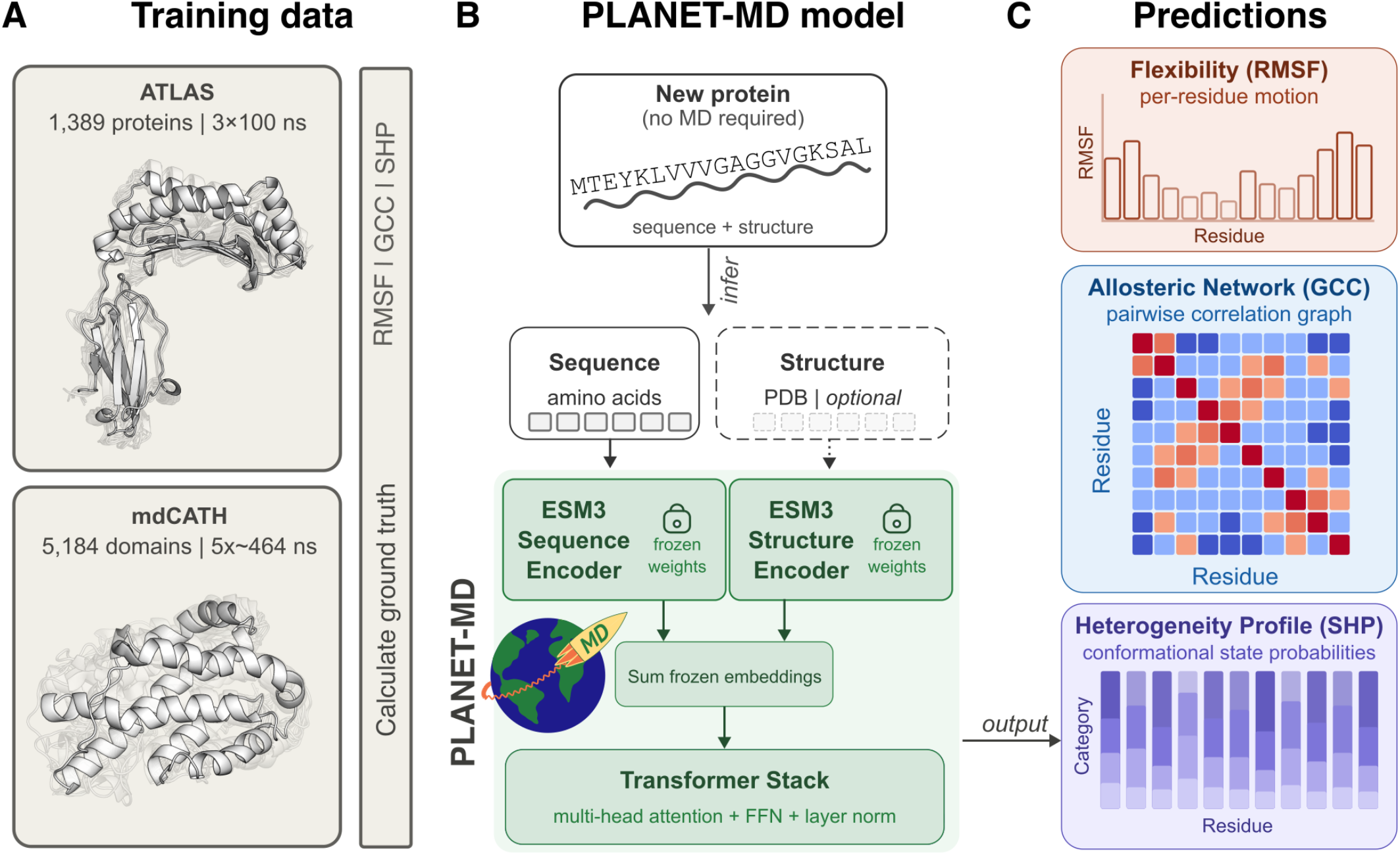
Overview of PLANET-MD. **(A)** We take advantage of newly released large-scale molecular dynamics (MD) data sets to learn the dynamic behavior of proteins. ATLAS (Schweke et al., 2023) contains 1,389 proteins with 3 replicates of 100ns MD simulation per system. mdCATH (Mirarchi, Giorgino, and De Fabritiis, 2024) contains 5,184 proteins with 5 replicates of ≈464ns per system. From these simulations, we compute ground truth properties that are used to supervise PLANET-MD training. **(B)** The model architecture uses frozen weights from the ESM3 (Hayes et al., 2024) sequence and structure encoders to generate embeddings, which are first summed and passed through a transformer stack, then into distinct output heads. Once trained, inference can be performed on a new protein without requiring MD simulation. Rather, only a sequence is required, and a static structure can optionally be provided to improve performance. PLANET-MD is capable of predicting three properties derived from MD simulations. The Root Mean Square Fluctuation (RMSF) is a per-residue scalar measure of local flexibility. The Gaussian Correlation Coefficient (GCC-LMI) is a pairwise measurement of correlated motion between residues, and can be treated as a network of allosteric communication. The Structure Heteroregenity Profile (SHP) is a per-residue categorical distribution over the states occupied by each residue, where each category is derived from a machine-learned tokenization of the local neighborhood around the residue (Methods).

## Results

### PLANET-MD accurately predicts dynamic protein properties

We find that PLANET-MD achieves highly accurate prediction of dynamic and allosteric properties. We trained PLANET-MD on a subset of the ATLAS data set, and evaluated property prediction on the test set (Methods). We compare to Dyna-1 (Wayment-Steele et al., 2025) and BioEmu (Lewis et al., 2025), two recently released machine learning methods for predicting protein dynamics. While Dyna-1 was designed to predict millisecond-scale motions rather than nanosecond fluctuations, to wanted to evaluate how it would perform at predicting across time scales. To facilitate a fair comparison, we calibrated its predictions using 100ns simulations on 6 proteins from RelaxDB (*RMSF* = 0.4067 × *Dyna* − 0.0224, Methods). This calibration step allows us to match Dyna-1 predictions more closely to the time-scale we evaluate. BioEmu is a sampling-based method, and we thus evaluated it with both 10 and 100 samples. We evaluate using multiple metrics: RMSE and Spearman *ρ* for flexibility predictions (RMSF), graph diffusion distance (GDD) (Hammond, Gur, and Johnson, 2013) and Ipsen-Mikhailov spectral distance (IMSD) (Jurman et al., 2011) for allosteric networks (GCC-LMI), and Kullback-Leibler divergence (Kullback, 1951) for structure heterogeneity profile (SHP) distributions (Methods). In addition, since efficient inference is a primary goal of PLANET-MD, we compare their relative run times. We note that competing models were not retrained, and so it is possible that some test set proteins appeared in their training. Nonetheless, PLANET-MD is both the fastest and the most accurate method across proteins of diverse CATH domains (Orengo et al., 1997) and sequence lengths.

Across all metrics, PLANET-MD achieves state-of-the-art performance (Table 1). We developed three model variants: our full sequence-structure model (PLANET-MD), a sequence-only model (PLANET-MD-seq), and a lightweight sequence-only model (PLANET-MD-mini). All variants of PLANET-MD require significantly less time for inference compared to competing methods; even the full model requires less than a tenth of a second per protein on average and is significantly faster than the similarly sequence-based Dyna-1. As BioEmu is a sampling-based method, it is substantially slower, even with the relatively small number of samples drawn (Figure 2A).

**Table 1:** PLANET-MD learns residue-level dynamics. We compare leading methods for predicting residue flexibility (RMSF), correlation (GCC-LMI), and structure heterogeneity (SHP) on the ATLAS data set. Lower is better for all methods except Spearman *ρ*. The best performance for each is **bolded**. All metrics are unitless, except RMSE which is in angstrom.

| Model | RMSF |  | GCC-LMI |  | SHP | Model Information |  |  |
| --- | --- | --- | --- | --- | --- | --- | --- | --- |
| | RMSE (Å) | Spear. $\rho$ (↑) | GDD | IMSD | KL-Div | Seq. | Struct. | Params. |
| PLANET-MD-mini | 1.04 | 0.680 | 1.023 | 2.532 | **1.089** | ✓ | ✗ | **1.5M** |
| PLANET-MD-seq | 0.90 | 0.719 | 0.944 | 2.387 | 1.225 | ✓ | ✗ | 29.7M |
| PLANET-MD | **0.83** | **0.789** | **0.859** | 1.898 | 1.551 | ✓ | ✓ | 30.2M |
| BioEmu (10 samples) | 1.92 | 0.715 | 2.550 | 3.357 | 1.923 | ✓ | ✗ | 31.2M |
| BioEmu (100 samples) | 2.02 | 0.748 | 1.215 | **1.871** | 1.778 | ✓ | ✗ | 31.2M |
| Dyna-1 | 3.61 | 0.267 | - | - | - | ✓ | ✓ | 188.9M |
| Dyna-1 (calibrated) | 1.91 | 0.267 | - | - | - | ✓ | ✓ | 188.9M |

**Figure 2:**
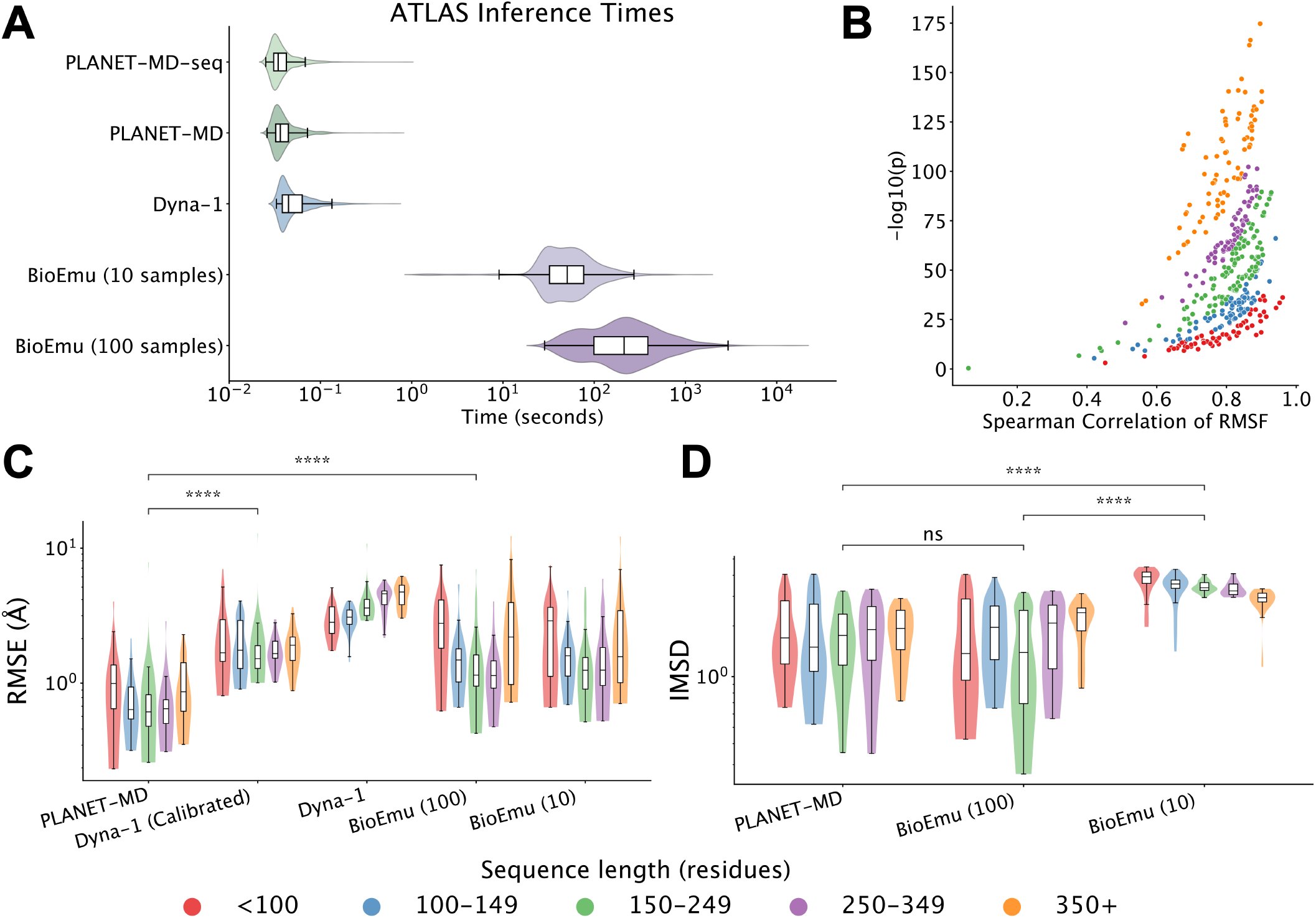
PLANET-MD predicts dynamic properties rapidly and accurately. **(A)** We show the distribution of per-protein inference times of each model on the ATLAS data set. PLANET-MD and its sequence-only variant are significantly faster than Dyna-1 (Wilcoxon signed-rank test, *p* = 5.61e − 72), requiring less than a tenth of a second per protein on average. BioEmu is substantially slower, requiring over a minute per protein, even with only ten samples. **(B)** We show the Spearman ρ of RMSF predictions plotted against significance of correlation (-log10(p value)) for each protein in ATLAS, colored by sequence length. PLANET-MD consistently achieves strong performance of flexibility, even for large proteins. **(C)** We compare the accuracy of RMSF prediction (root mean square error, angstom) on the ATLAS data set between PLANET-MD, Dyna-1 (and a calibrated version), and BioEmu (with 10, 100 samples). PLANET-MD achieves the most accurate prediction of RMSF, significantly better than both Dyna-1 (Mann-Whitney U test, *p* = 1.10*e* − 29) and BioEmu (*p* = 1.82*e* − 19). **(D)** We also compare the accuracy of GCC-LMI prediction, evaluated using the Ipsen-Mikhailov Spectral Distance (ISMD) as a measure of graph similarity. We compare only with BioEmu, as Dyna-1 can not predict allosteric networks. PLANET-MD outperforms BioEmu with 10 samples (*p* = 1.23*e* − 32), and achieves competitive performance with BioEmu with 100 samples (*p* = 0.74).

For flexibility prediction, PLANET-MD is the most accurate model (*RMSE* = 0.83, Spearman ρ = 0.789). Crucially, PLANET-MD maintains strong performance even when applied to very large proteins. While most proteins analyzed are between 100-350 amino acids, the longest proteins can be upward of 1500 amino acids, a scale which significantly challenges current methods. PLANET-MD maintains high accuracy across this range with significant correlation to ground truth values (Figure 2B). PLANET-MD significantly outperforms both Dyna-1 (*p* = 1.10e − 29) and BioEmu (*p* = 1.82e − 19); When performance is broken down by sequence length, we observe with Dyna-1 struggling particularly on longer proteins and BioEmu showing reduced accuracy at both extremes (Figure 2C).

PLANET-MD also reconstructs highly accurate allosteric networks, or par with the slower BioEmu. When measured by GDD, PLANET-MD is the most accurate (0.859 vs. 1.215), while BioEmu outperforms PLANET-MD in IMSD (1.871 vs. 1.898). In neither case is this difference statistically significant. Thus, we find PLANET-MD to be a strong option for investigating these networks at scale without the need for explicit ensemble sampling via ML-based samplers or molecular dynamics. Below, we show a case study probing the allosteric network of KRAS (Figure 6). We likewise find that directly predicting a distribution over structure classes achieves competitive accuracy with explicit ensemble sampling. Measured by the KL divergence to the true distribution, PLANET-MD SHPs are more accurate than those from BioEmu (1.551 vs. 1.778); in fact the most accurate distributions were predicted by the mini variant (1.089 KL-div). These results support the conclusions from Kalifa, Horvitz, and Radinsky, 2026, where they found that PLANET-MD SHP conditioning was sufficient to learn state-aware representations in DynamicsPLM.

### Dense structure features improve prediction accuracy

We next performed an ablation study of structure input representations for PLANET-MD. Figure 3A illustrates the ESM3 VQ-VAE structure encoder (Hayes et al., 2024) with its geometric attention transformer (green, *d* = 1024), projection layer (orange, *d* = 128), and quantization (blue, *d* = 1, 4096 categories). We compared these representations against a sequence-only model (gray, zero-vector structure embedding) and Ramachandran angles (*ϕ, ψ*) as baseline structure features (purple, *d* = 2). Validation loss curves show dense structure features outperform baselines, but quantized representations perform similarly to simple Ramachandran angles (Figure 3B). Notably, the sequence-only model performs comparably to basic structure representations, motivating its development for broader applicability. We compare full the PLANET-MD versus PLANET-MD-seq on test proteins by measuring RMSF RMSE, GCC-LMI IMSD, and SHP KL Divergence (Figures 3C-E). The structure-based model consistently outperforms the sequence-only variant, motivating the inclusion of structure features for inference. However, the sequence-only model still maintains accurate performance, and can be used for extremely high-throughput inference.

**Figure 3:**
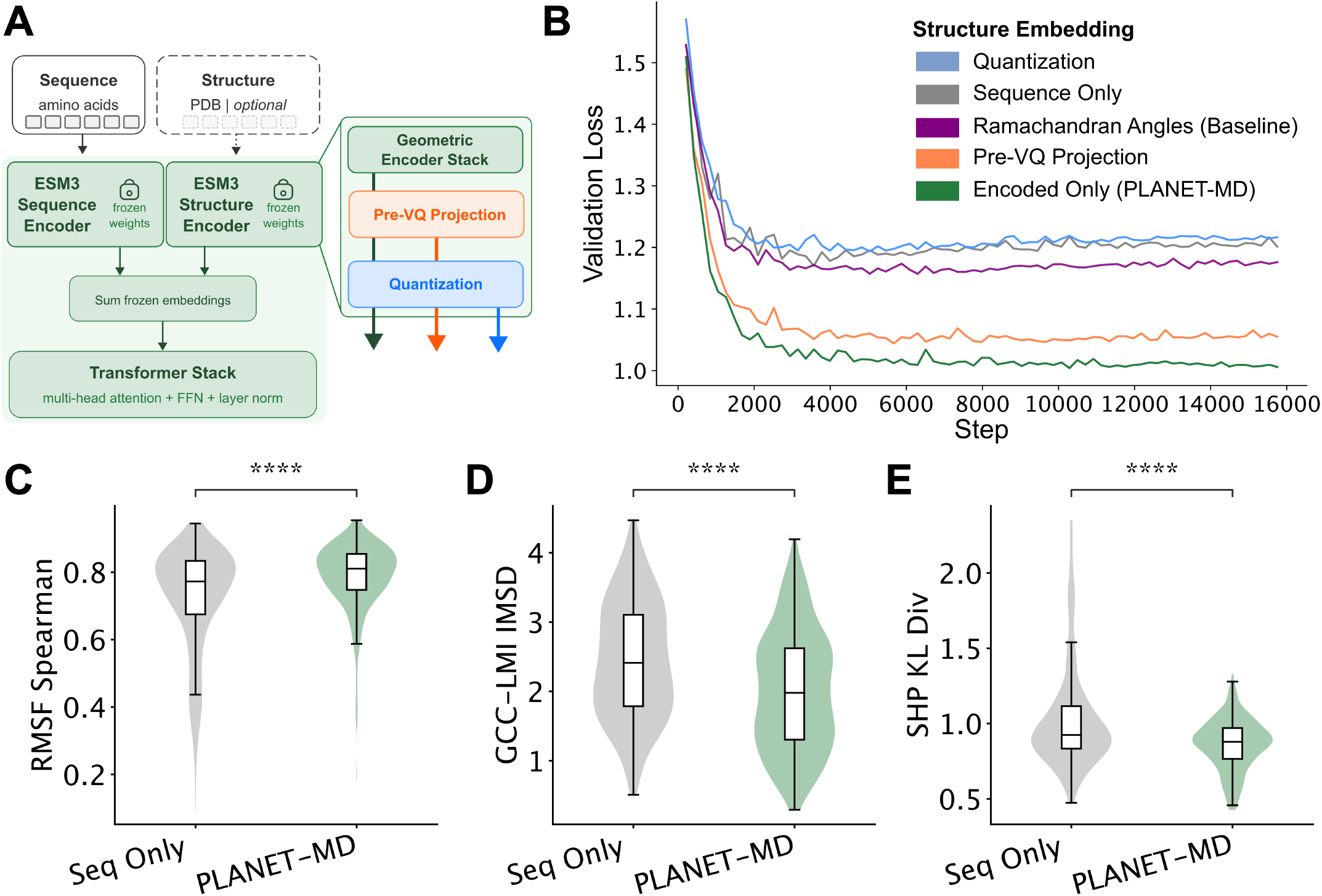
Evaluating the impact of diffierent structure representations. **(A)** The ESM3 structure encoder is used with frozen weights to generate features for PLANET-MD. However, this encoder actually contains several steps, and we sought to evaluate which representation would achieve the best performance. The primary output is quantized structure tokens (blue), but we evaluate also the projected vectors immediately prior to quantization (orange), and the embeddings from the geometric encoder (green). We compare these with a baseline structure embedding, using the Ramachandran angles for each residue (purple), and with a sequence-only model (gray, zero vectors for structure). **(B)** We train a model with each set of structure features, and show the validation loss curves for each method. Dense structure features (encoded, projected) outperform baseline, sequence-only, and quantized structure features. Based on this ablation study, we selected to use the encoded structure features for the full PLANET-MD model. **(C)** We compare the full PLANET-MD model to the version using only sequence features, and find significant performance gains from including static structures. The full model achieves higher Spearman correlations for RMSF (Mann-Whitney U test, p = 2.06e − 9), **(D)** lower IMSD for allosteric network prediction (p = 8.71e − 12), and **(E)** a lower KL divergence to the true structure heterogeneity profile (p = 2.73e − 11). However, the sequence-only model still remains broadly accurate, motivating our development of PLANET-MD mini.

### Validating PLANET-MD with experimental relaxation data

In recent work, Wayment-Steele et al., 2025 released RelaxDB, a database of NMR relaxation measurements for 133 proteins. They use *R*_*ex*_ values to validate Dyna-1 estimates of *μs* − *ms* motions, but this rich data set also contains measurements of heteronuclear NOE values (hetNOE), which measures the type of *ps* − *ns* that we expect PLANET-MD to capture (Figure 4A). We observe that PLANET-MD RMSF is a zero-shot predictor of hetNOE; in the 133 predictions in RelaxDB, hetNOE is highly anticorrelated with predicted RMSF (Figure 4B, *n* = 8249, ρ = −0.67). Since lower hetNOE indicates more flexible motion, we use ReLU(1 − *RMSF*) to predict directly; this achieves an AUPR of 0.95, and an accuracy of 0.87 with the threshold of predicted hetNOE < 0.65 established by Wayment-Steele et al., 2025 (dashed gray lines). We highlight one case study, the Cas9 inhibitor ACRIIA4 highlighted in the initial RelaxDB manuscript. We find a high level of concordance between the experimentally determined hetNOE and the PLANET-MD RMSF predictions (Figure 4C,D), validating our approach as an accurate predictor of local fluctuations.

**Figure 4:**
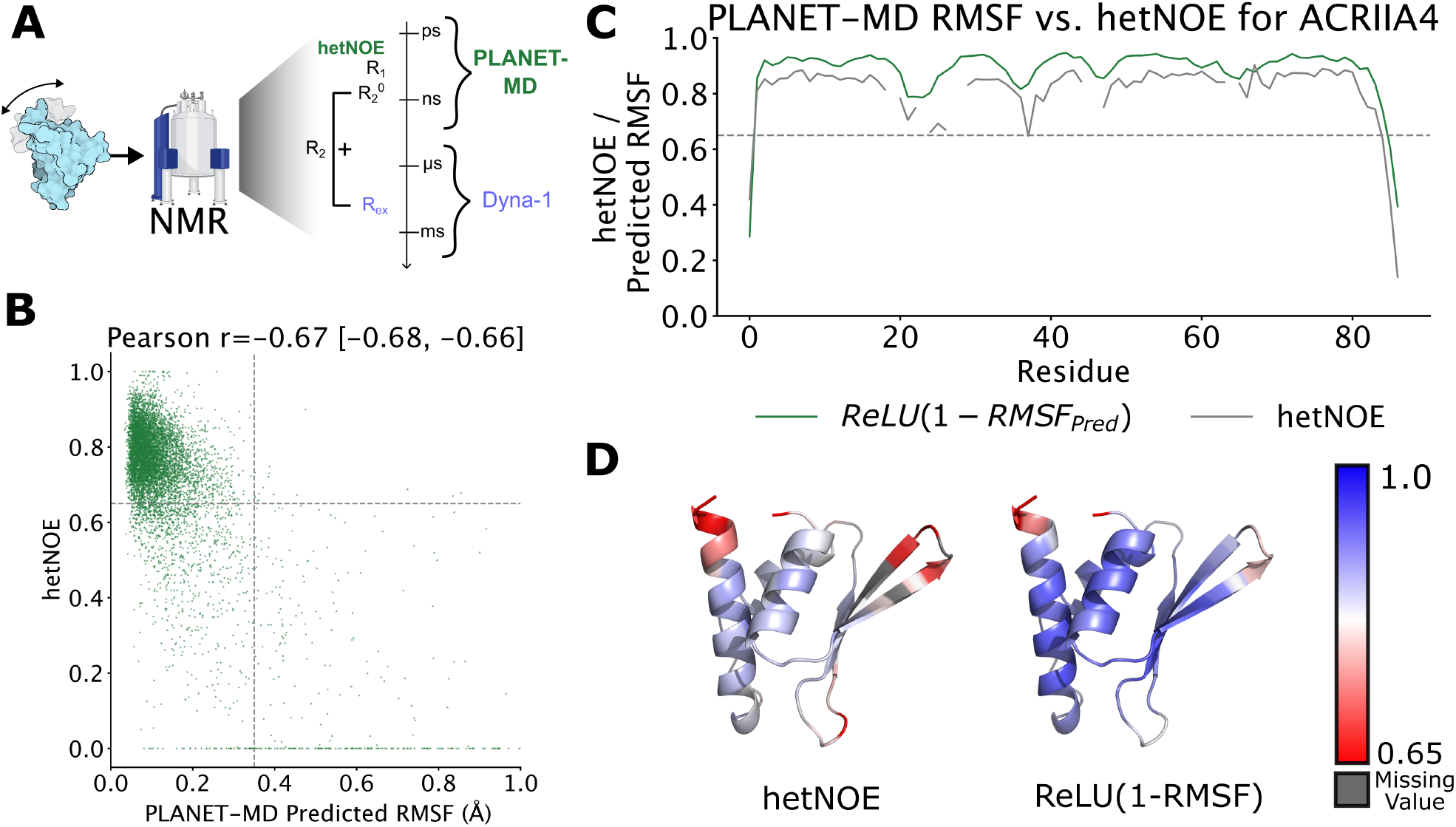
Validating PLANET-MD with experimental hetNOE measurements. **(A)** Nuclear magnetic resonance (NMR) experiments report several measurements of molecular motion corresponding to several time scales; while Dyna-1 focuses on missing *R*_*ex*_ values corresponding to slower movements, PLANET-MD is trained on shorter simulations and is thus more likely to predict the faster motions captured by hetNOE. **(B)** RelaxDB contains curated NMR hetNOE measurements for 133 proteins. We show the predicted RMSF for each residue (*n* = 8, 249), with a dashed gray line at 0.65, the reported threshold for *ps* − *ns* motion in Wayment-Steele et al., 2025. Predicted RMSF is highly predictive of experimental hetNOE (Pearson r = − 0.67, AUPR= 0.95, binary accuracy = 0.87). **(C)** We show example data for one protein from RelaxDB, the Cas9 inhibitor ACRIIA4. We compute ReLU(1− *RMSF*) to account for the fact that a lower hetNOE corresponds with more flexible motion, or a higher RMSF. **(D)** We show the true and predicted flexibility values on the structure of ACRIIA4, accurately identifying regions of the protein with *ps* − *ns* motion. Here, a lower value (< 0.65) corresponds to motion at this time scale.

## Modeling flexibility as a function of temperature

PLANET-MD is the only one of the three methods studied here that is able to take temperature as input. Since the mdCATH data set contains simulations of the same protein at five temperatures, we next evaluate the ability of PLANET-MD to scale RMSF prediction across temperature. We compare to a baseline from Sá Ribeiro and Lima, 2023, using the relation *B*_*obs*_ = *B*_0_*e*^*kT*^ to scale the RMSF at 320K to other temperatures (Methods). While PLANET-MD performance is comparable with this baseline at lower temperatures, at much higher temperatures we significantly outperform the naive scaling approach, as different residues will have different responses to temperature and a single scale across all residues is likely to collapse this differing response (Figure 5A,B). We show one case study the *E. coli* glutaminase *ybaS* that is differentially affected by increased temperatures (Figure 5C). While some regions remain stable, others experience significantly increased fluctuations and even unfolding. We find that PLANET-MD maintains strong predictive performance below 400K, but starts to become less accurate at higher temperatures (Figure 5D). In this regime, RMSF has higher variability across any given slice of simulation because the protein explores a larger conformational space (Discussion). Comparisons with BioEmu and Dyna-1 using only the lowest temperature of mdCATH simulations (320K) are provided in Table A1 (Appendix A.2).

**Figure 5:**
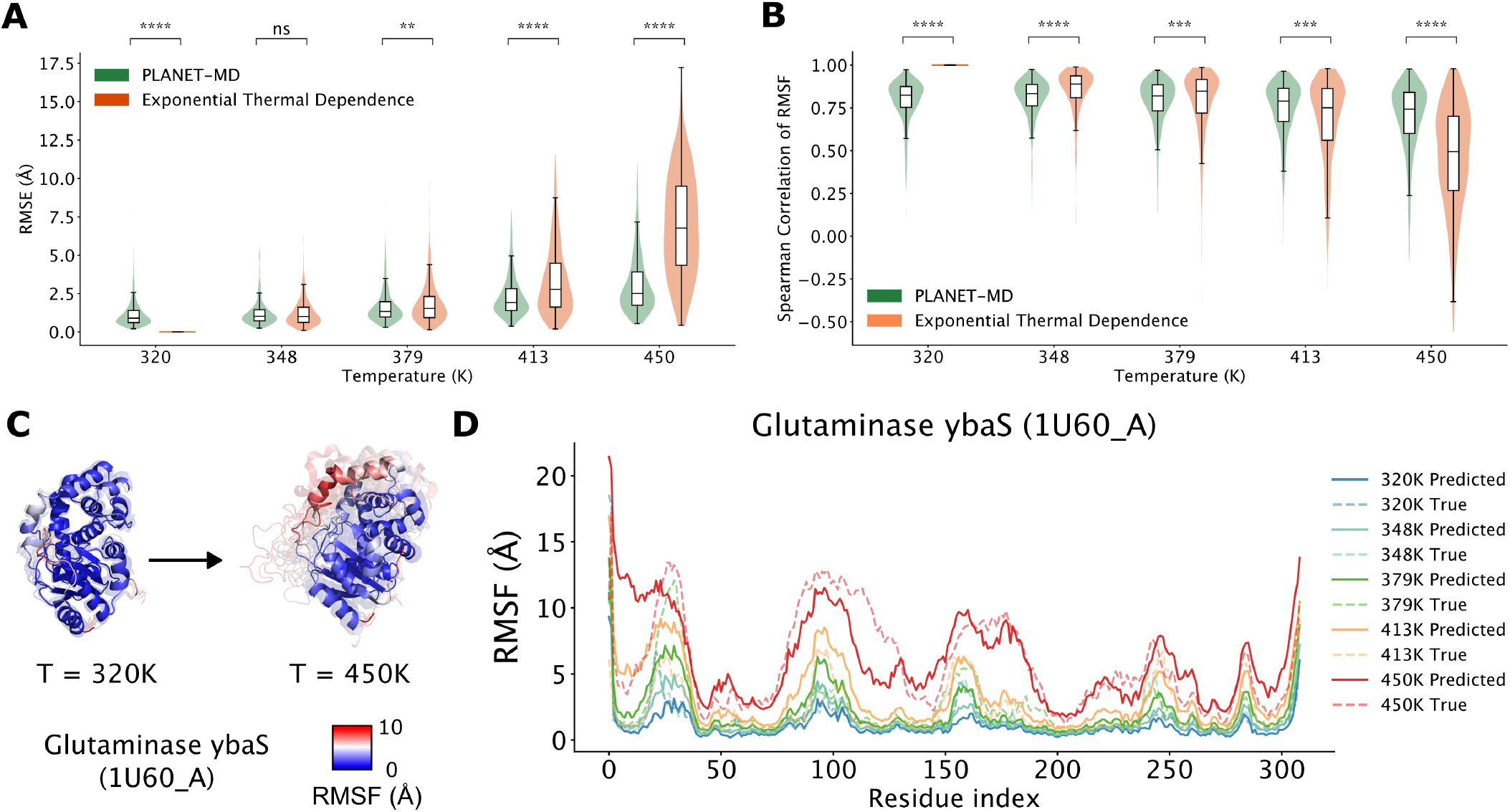
PLANET-MD accurately predicts the impact of temperature on flexibility. **(A)** We show the PLANET-MD root mean squared error (RMSE) of RMSF across temperatures for proteins from the mdCATH data set, compared to a baseline model that scales the lowest temperature exponentially (Sá Ribeiro and Lima, 2023). This baseline has by definition zero error at 320 Kelvin (K), the base temperature that is then scaled to higher temperatures. PLANET-MD consistently predicts RMSF accurately than this exponential scaling model, demonstrating the value of explicitly conditioning on temperature in model training and inference. **(B)** We report also the Spearman correlation of RMSF for both methods, finding similar results. PLANET-MD is especially strong compared to the baseline at the highest temperatures, suggesting that the exponential relationship breaks down with larger gaps between temperature. **(C)** We show example structural ensembles for one case study protein, the *E. coli* glutaminase *ybaS* (PDB: 1U60 chain A). These structures were sampled from molecular dynamics simulations. We show here 10 sample structures from simulations at 320K and at 450K, colored by RMSF. As temperature increases, some regions of *ybaS* stay rigid, while others experience significantly increased flexibility and even occasional unfolding. **(D)** PLANET-MD accurately predicts RMSF across five different temperatures for *ybaS*. We show both the true (dashed) and predicted (solid) RMSF, colored by temperature. As observed in the aggregate mdCATH data, RMSF becomes more difficult to predict accurately at the highest temperatures (413K, 450K).

### Building an allosteric network in KRAS

We designed PLANET-MD to enable building and interpreting allosteric networks. To evaluate this capability, we turn to the protein *KRAS*. Mutations in *KRAS* are widely implicated in cancers such as colorectal (Yamauchi et al., 2012), lung (Suda, Tomizawa, and Mitsudomi, 2010), and pancreatic (Leroux and Konstantinidou, 2021). Thus, *KRAS* is a popular candidate target for therapeutic development (Cox et al., 2014; Ryan and Corcoran, 2018; Wolpin et al., 2026). However, *KRAS* displays highly dynamic behavior, complicating the development of effective drugs. It acts as a “molecular switch,” only relaying signals when bound by GTP, the binding of which induces a conformational change in the structure. We thus applied PLANET-MD to explore conformational heterogeneity in *KRAS*. We first validated PLANET-MD predictions of RMSF, GCC-LMI, and SHP for *KRAS* compared to ground truth from an in-house simulation (Appendix A.3). Then, following Maschietto et al., 2023, we constructed a network from the predicted GCC-LMI by applying a distance mask (using the predicted *Cα* distances) and treating pairwise correlations as weighted edges (Figure 6A). We next applied a Girvan-Newman clustering algorithm to hierarchically divide the network into communities (Figure 6B, *k* = 5). Distinct structural regions emerge, including the structural core (gray), the disordered C-terminal tail (yellow), a ligand binding pocket (blue, PDB:4DSN, Maurer et al., 2012), and a pair of anti-parallel helices (green) (Figure 6C). Interestingly, the arginine at position 73, part of a helix turn, clusters entirely on its own. We compare these unsupervised clusters with known region annotations in KRAS (Pantsar, 2020), and find that the ligand binding cluster corresponds strongly with the Switch I region, the anti-parallel helices with Switch II, and the C-terminal tail with the highly variable region (HVR) (Figure 6F).

**Figure 6:**
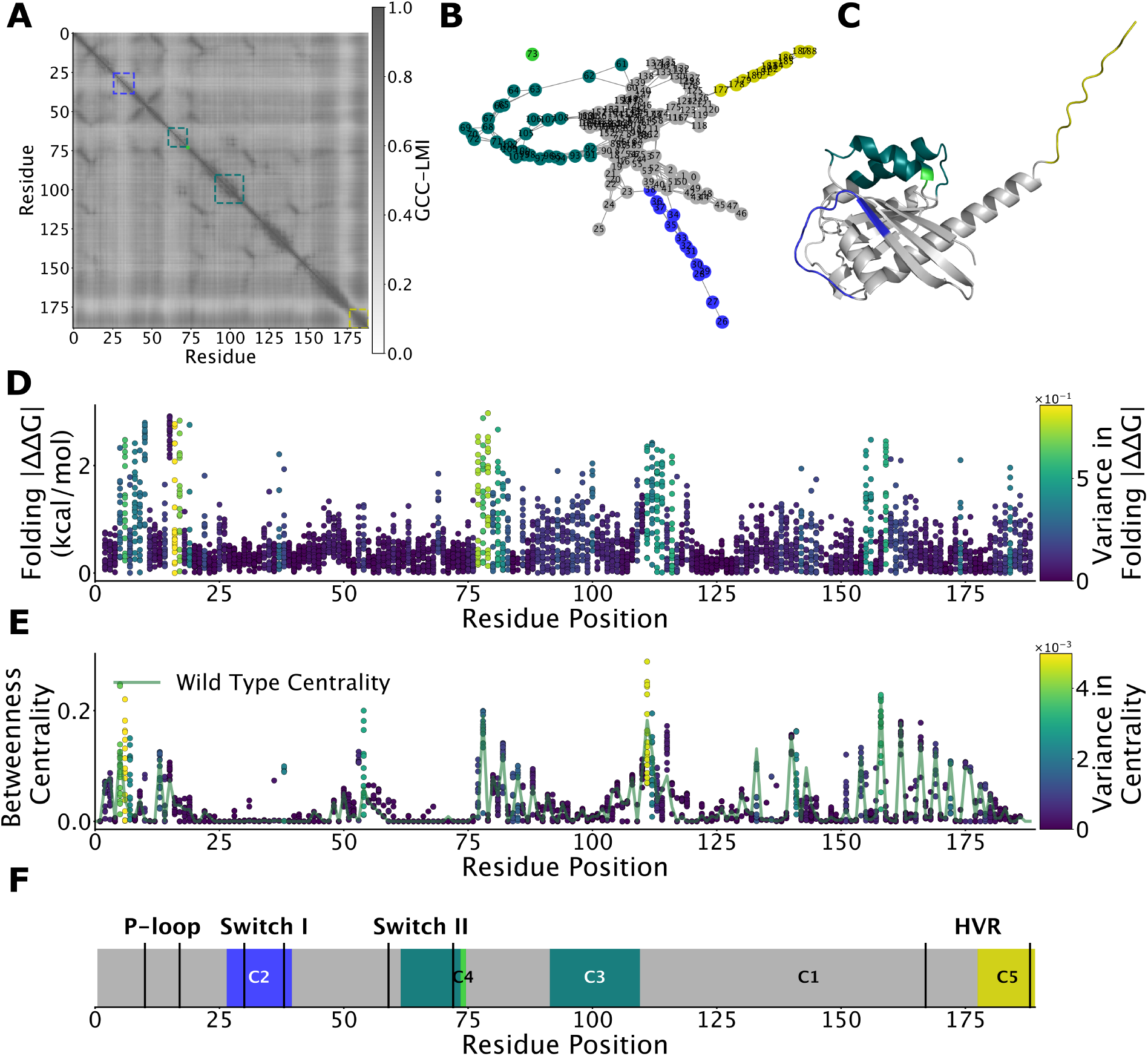
Mutations in allosteric networks recapitulate folding free energy in KRAS. **(A)** We show the pairwise correlations (GCC-LMI) in per-residue motion predicted by PLANET-MD, which we interpret as edge weights in a network (edges are first filtered by *Cα* distance ≤8). Overlaid on this heatmap are four clusters identified by unsupervised community detection (Girvan-Newman clustering). **(B)** The predicted allosteric network, colored by community. We color the largest community gray, as this acts as a “leftover” cluster. **(C)** When network clusters are projected onto the structure of KRAS, we find distinct structural modules, including a structural core (gray), binding pocket/Switch I (blue), Switch II (teal), and 2024 the hypervariable C-terminal tail (yellow). **(D)** Weng et al., perform a deep mutational scan (DMS) of KRAS, measuring the ΔΔ*G* for each possible mutant at each location. We show the results of that DMS here, colored by the variance at each position. **(E)** We perform an *in silico* DMS with PLANET-MD, reporting first the wild type betweenness centrality of each residue in the network (green), as well as the betweenness centrality of that residue when mutated. Points are colored by variance in centrality at each position. We find that both the wild type centrality and the predicted variance correspond with variance in Δ Δ*G* of folding for *KRAS*. Thus, this approach is a meaningful computational option for identifying functionally meaningful residues in KRAS. **(F)** We show the PLANET-MD predicted clusters, overlaid with annotated regions determined by Pantsar, 2020. While the P-loop is not identified in a cluster, PLANET-MD clusters otherwise correspond to well-documented structural regions, including Switch I (blue), Switch II (teal), and the C-terminal highly variable region (yellow). Switch I is known to be a location of ligand binding to KRAS (PDB: 4DSN).

Finally, we sought to apply this network model to explore residue-level function in KRAS. Weng et al., 2024 performed a deep mutational scan (DMS) of *KRAS*, measuring the change in free energy (Δ Δ*G*) of folding (Figure 6D) and with several different binding partners. Maschietto et al., 2023 then showed that allosteric networks derived from molecular dynamics simulations could be used to recover the results of this study. We take this previous study further: because betweenness centrality measures nodes with key roles in information flow in the network, we hypothesized that residues which control dynamic structure in the allosteric network would have high variability in betweenness centrality when mutated. We find that allosteric networks predicted by an *in silico* DMS with PLANET-MD can similarly recapitulate this experimental study in less than an hour of GPU time, identifying functionally important residues in *KRAS* (Figure 6E). The scalability of PLANET-MD enables this level of functional characterization of allosterically important residues at the scale of the proteome.

### Proteome-wide study of structure dynamics

The scalability of PLANET-MD also enables proteome-wide analyses unavailable to more traditional simulation methods or even ML-based samplers like BioEmu (Figure 7A). Thus, we applied PLANET-MD to analyze how flexibility is distributed across the human proteome. We download over 20,000 human protein structures from AlphaFoldDB (version 4) (Varadi et al., 2022), and obtain domain/region annotations and subcellular locations for each protein from UniProt. We use PLANET-MD to predict RMSF, GCC-LMI, and SHPs for all proteins, which took just over 73 minutes on a single A100 GPU (including reading and writing from disk). Unsurprisingly, we find that residues annotated as part of a disordered region have a significantly higher predicted RMSF than residues annotated as part of an ordered region (Figure 7B, Mann-Whitney U test, p < 10e − 300, underflow float64 precision). We next sought to evaluate the distribution of flexible residues across different subcellular compartments, and whether proteins localized to a particular compartment are more likely to contain flexible regions. While most human proteins are cytoplasmic (30.6%) or nuclear (28.8%), there are also a substantial number that are membrane associated (22.2%), secreted (12.6%), or mitochondrial (3.1%) (Figure 7C). We find that nuclear proteins tend to be the most flexible, and are significantly more so than proteins from other compartments (Figure 7D). However, we note that there is a high level of variance in flexibility in all compartments, and that subcellular location is not the primary driver of variance in flexibility.

**Figure 7:**
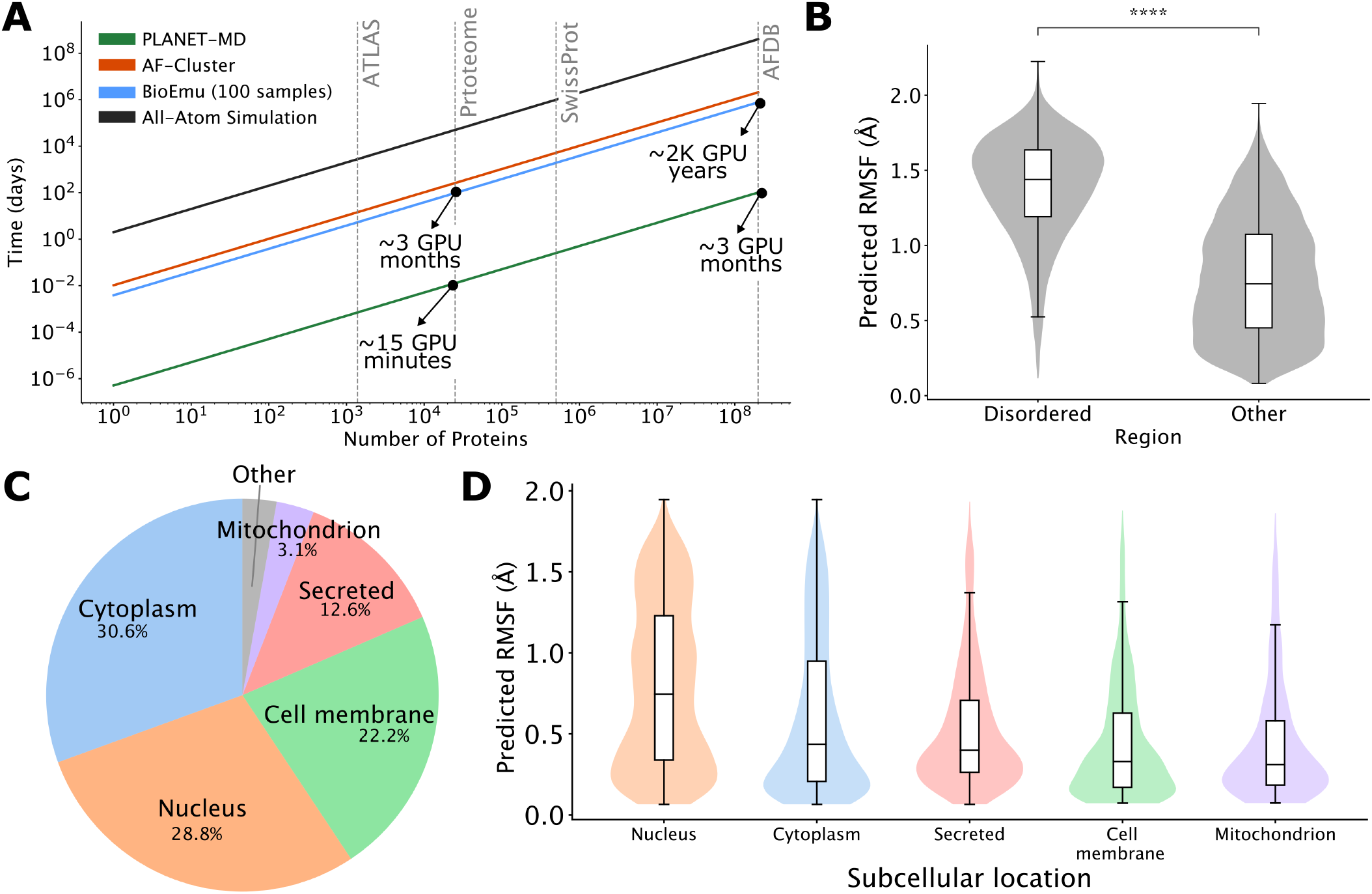
PLANET-MD enables the first proteome-scale prediction of human protein dynamics. **(A)** Due to the combination of rapid and accurate inference, PLANET-MD is well suited to scale prediction of protein dynamics to large sequence corpora. We show here estimated runtimes for PLANET-MD (green), AF-Cluster (Wayment-Steele et al., 2024) (orange), BioEmu (Lewis et al., 2025) (blue), and all-atom MD using OpenMM (Eastman et al., 2023) (black). To run inference on all of the AlphaFold Database (Varadi et al., 2022) using AF-Cluster or BioEmu would take approximately 2,000 GPU years; PLANET-MD could perform similar inference in ≈ 3 GPU months. To demonstrate the capabilities enabled by this throughput, we perform an analysis of the dynamics of the human proteome, which takes roughly 15 minutes on a single GPU with PLANET-MD. **(B)** Using sequence region annotations from SwissProt, we compare the predicted RMSF of residues labeled as disordered with other residues. We find these residues to have significantly higher predicted RMSF (Mann-Whitney U test, *p* < 10*e* −300) than ordered amino acids. **(C)** We show the relative proportion of sub-cellular locations for human proteins annotated in SwissProt. Most proteins are cytoplasmic (30.6%) or nuclear (28.8%), while fewer are localized to the cell membrane (22.2%), secreted (12.6%), or mitochondrial (3.1%). **(D)** Using PLANET-MD predicted RMSF, we compare the distributions of flexibility for proteins from each compartment. On average, nuclear proteins tend to be more flexible than those from other compartments.

## Discussion

In this paper, we introduce PLANET-MD, a method for ultra-fast modeling of protein dynamics and allosteric networks. Leveraging protein foundation models and MD simulations, PLANET-MD enables proteome-scale analysis while maintaining accuracy across diverse protein families, structures, and lengths. Our approach accurately predicts RMSF and GCC-LMI, and introduces structure heterogeneity profiles (SHPs) as a novel representation of protein dynamics, providing a scalable intermediate-resolution view of a protein’s conformational ensemble. While we compare with Dyna-1 (Wayment-Steele et al., 2025), we view these models as complementary—capturing dynamics across different time scales requires multiple approaches. One limitation of PLANET-MD is its training on relatively short MD trajectories (100-500ns). While short time-scale fluctuations are able to capture local allosteric networks (Singh and Bowman, 2017), these shorter simulations limit its ability to capture major conformational shifts that occur at longer timescales or higher temperatures, as suggested by the decreased performance on mdCATH at above 400K. As MD simulation data improves in quantity and quality, PLANET-MD will scale accordingly, but currently rare conformational states or correlations may be missed. Longer and more diverse simulations will also require additional data processing such as clustering trajectories by RMSD to maintain fidelity of local metrics like RMSF. Importantly, PLANET-MD predicts dynamic properties rather than generating structures or sampling from the Boltzmann distribution, limiting its usage in cases where a full ensemble is required. However, work from Kalifa, Horvitz, and Radinsky, 2026 suggests a path forward for using SHPs to explicitly sample ensembles, which we will explore in future work. Additionally, our reliance on ESM3 (Hayes et al., 2024) and other pre-trained PLMs means performance may degrade for proteins less constrained by evolution, such as antibody hypervariable regions or intrinsically disordered proteins. Finally, to create allosteric networks we selected the generalized correlation coefficients with linear mutual information (GCC-LMI) as implemented in MDiGest (Maschietto et al., 2023), which was useful for its speed in analyzing a large number of individual trajectories. There exist a variety of different ways that one can calculate this connectivity (Bernetti et al., 2024), which will be further explored in future variants of PLANET-MD. Despite these limitations, PLANET-MD is poised to have an impact in systems and structural biology. By dramatically reducing computational requirements for protein dynamics research, PLANET-MD democratizes access to this crucial area, helping to bridge the gap between well-funded research groups and smaller labs or companies. This reduced computational footprint comes with environmental benefits, decreasing energy consumption for dynamics studies by orders of magnitude. This scale also opens the door for further genome-wide analyses, allowing researchers to study the link between variant pathogenicity and allosteric network centrality, or to probe the effects of point mutations on protein flexibility. We are releasing PLANET-MD with open, permissive licensing to enable widespread use within the research community at https://github.com/flatironinstitute/PLANET-MD.

## Acknowledgments

The authors would like to thank Siavash Golkar, Miro Astore, Pilar Cossio, and Abhilash Sahoo for helpful discussions regarding this work. We especially thank Federica Maschietto for extremely helpful guidance on the *KRAS* deep mutational scanning experiments and the use of MDigest to compute correlated fluctuations. S.S. and S.M.H. are supported by the Simons Foundation. The Flatiron Institute is a division of the Simons Foundation. S.M.H. does not have any competing interests to declare. S.S is an employee and shareholder of Divergence Labs.

## Methods

### Simulation Data

We collected training data from two publicly-available repositories of MD simulations, ATLAS (Schweke et al., 2023) and mdCATH (Mirarchi, Giorgino, and De Fabritiis, 2024). ATLAS contains 1,389 single-domain protein chains, each simulated for three replicates of 100 nanoseconds at 300K. Simulations were performed with GROMACS v2019.4 with the CHARMM36m force field (Van Der Spoel et al., 2005; Huang et al., 2017). mdCATH contains 5,184 protein domains, each simulated for five replicates of average length 464 nanoseconds at five different temperatures (320K, 348K, 379K, 413K, 450K). Simulations were performed with ACEMD on GPUGRID.net (Harvey, Giupponi, and Fabritiis, 2009). Both ATLAS and mdCATH selected chains from diverse CATH domains (Knudsen and Wiuf, 2010), but we perform an even more rigorous structural similarity-based split using FoldSeek (Kempen et al., 2024) with a 30% similarity threshold. We split clusters into training (80%), validation (10%), and test (10%) and assign all proteins sharing a cluster to the same split. This split strategy is crucial for properly evaluating generalization to unseen protein structures, as it tests the model’s ability to predict dynamics for structural motifs it has not encountered during training.

### Computing derivatives

Using the simulation data, we compute three information-dense derivatives that capture conformational diversity and allosteric communication in the protein. A protein can be represented both by its sequence of amino acids *X* ∈ {1, …, 20}^*N*^ = *x*_1_, …*x*_*N*_ and by its structure *S*^0^ ∈ ℝ^*N ×*3^, containing the positions (three spatial coordinates) of each alpha carbon (*Cα*). We define a simulation trajectory *S* ∈ ℝ^*N ×t×*3^ containing these positions simulated for *t* time steps. Although the full simulations contain all-atom positions, we focus here on only *Cα* motions and filter the trajectory accordingly. Prior to downstream computations, the trajectories are centered and aligned using mdtraj `center_coordinates` (McGibbon et al., 2015).

### Residue fluctuations

The root mean squared fluctuation (RMSF) is a scalar measure of the flexibility of a residue over the course of the simulation. For each residue *x*_*i*_ ∈ *X* we compute the RMSF as

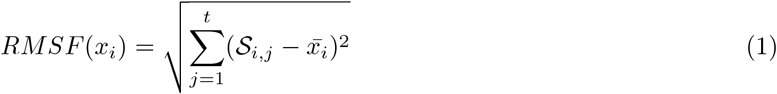

with 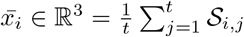We use the RMSF computation as implemented in mdtraj (McGibbon et al., 2015), v1.10.2.

### Pairwise correlations

To capture the effects of residue communications, we compute a measure of pairwise correlated motion using the Gaussian correlation co-efficient with linear mutual information (GCC-LMI). The GCC-LMI is a powerful yet scalable way to capture pairwise motions within the trajectory, although it is limited to capture only linear correlations. We choose the linear mutual information over the full GCC because of its improved scaling for large proteins and trajectories, and because we expect the most immediate signals of residue correlation to be linear. We compute GCC-LMI as

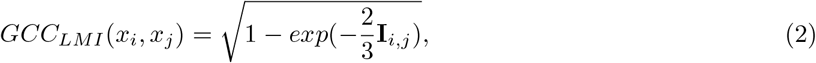

where the linearized mutual information between *x*_*i*_ and *x*_*j*_ can be estimated by a Gaussian estimator as

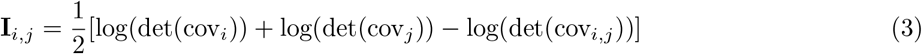

with

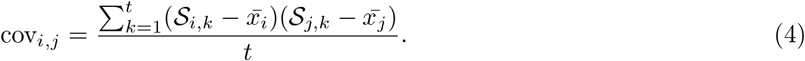

We use the GCC-LMI computation as implemented in MDigest (Maschietto et al., 2023), v0.1.8.

### Tokenized structure heterogeneity profiles

Finally, we calculate a structure heterogeneity profile (SHP) to capture the distribution of states that a residue visits during the simulation–while these simulations are likely too short to capture the full ensemble of states, the SHP gives a higher dimensional view into the diversity of protein conformations, and allows us to supplement the linear measure of flexibility (RMSF) with a categorical measure of residue shape space. To compute the SHP, we use the FoldSeek structure tokenizer (Kempen et al., 2024). Briefly, FoldSeek generates a sequence *FS*(*S*^0^) ∈ 3*DI*^*N*^, where the set 3*DI* is an alphabet defined by training a vector-quantized variational autoencoder (VQ-VAE) to reconstruct protein structures (|3*DI*| = 20). This linear encoding of the 3-dimensional protein structure has proven useful for structural similarity search, and each token in the alphabet represents a discrete region of backbone structural space.

We then compute

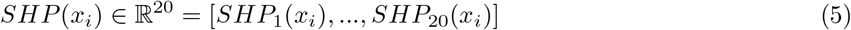

with 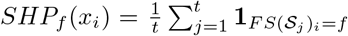. For computational efficiency, and after making the observation that the distribution *SHP*(*x*_*i*_) is relatively stable over different subsets of the simulation, we compute this distribution not over every frame but over every *k*^*th*^ frame of the trajectory.

### Model

#### Pretrained Model Features

We leverage the foundation model ESM3 (Hayes et al., 2024) to generate initial representations of both protein sequence and structure (Figure 1). The parameters of the ESM3 model are frozen and not updated during training. For the full model, we use the “encoded” output of the ESM3 structure VQ-VAE. For the sequence-only version of the model, we use a matrix of all zeros for the structure encoding. We define *S*^0^ ∈ R^*N ×*3^ as the known structure for a protein; for all training data we take the first frame of the simulation as this *a priori* known structure. This joint encoding is then passed through a transformer encoder *TE* to generate a shared hidden representation *H*(*X, S*^0^).

#### Classification Heads

The transformer encoder is defined as

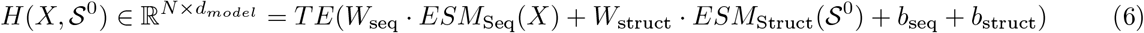

Where 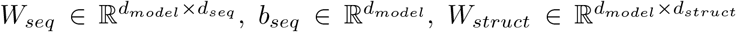 and 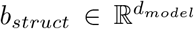 are the learnable parameters of the linear layers applied to the sequence and structure representations, respectively. This hidden representation is then passed into each classification head. The temperature *T* is first concatenated along the feature dimension for RMSF prediction. This is used to predict temperature-dependent flexibility on the mdCATH data set. This representation, without temperature included, is also used to predict the SHP.

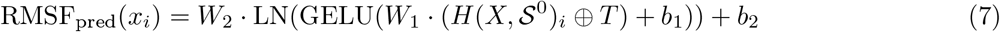

where LN refers to layer normalization and ⊕ represents feature-dimension concatenation.

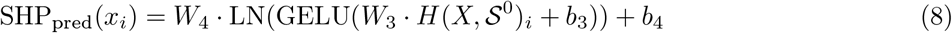

The two pairwise predictions, *GCC*_*LMI− pred*_ and *CA*_*Dist* use a pairwise representation *Z*_*i,j*_ = (*H*(*X, S*^0^)_*i*_*· H*(*X, S*^0^)_*j*_) as input.

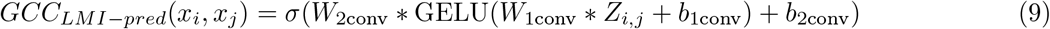

where * refers to the 2D *k*-convolution.

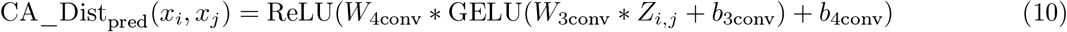

### Training

#### Losses

We compute a loss for each derivative predicted by the model, including the alpha carbon distances. The total loss is a weighted sum of these individual losses:

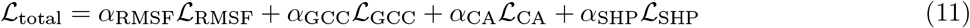

For the RMSF, GCC-LMI, and *Cα* distance predictions, we use the mean squared error (MSE) loss, and for the SHP predictions we use the Kullback-Leibler divergence loss, which is suitable for comparing probability distributions. We use *α* = 1 for all except *α*_*SHP*_ = 0.1 to balance the relative magnitude of each loss. We train with a batch size of 1 and gradient accumulation over 16 steps for an effective batch size of 16. We minimize the loss using the Adam optimizer with learning rate 5*e*− 5.

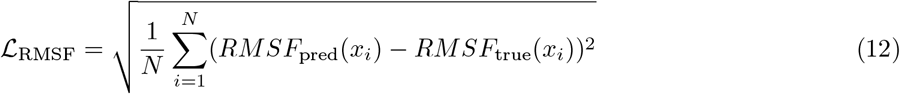

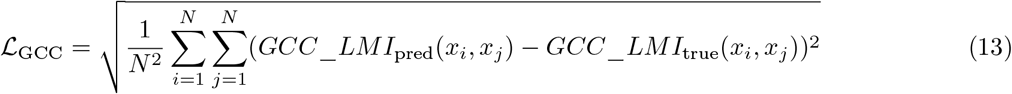

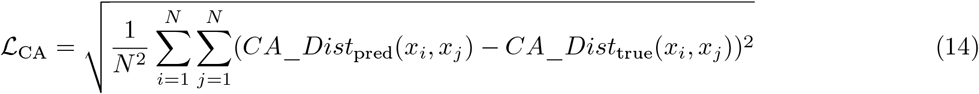

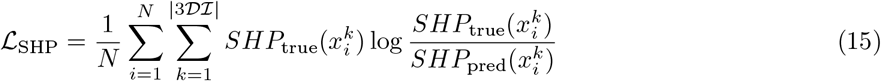

#### Performance Metrics

For measuring the quality of residue fluctuation predictions (RMSF), we use the root mean square error (RMSE) and Spearman correlation (ρ). Given *N* matched predictions *Y* with true labels *ŷ* we compute

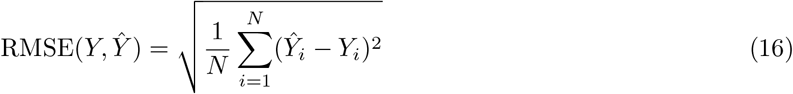

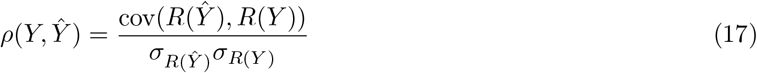

where *R*(*X*) is the rank ordering of *X* and *σ* (*·*) is the standard deviation. Since we are interpreting the GCC-LMI as allosteric networks, we use two graph-based distance metrics, the graph diffusion distance (GDD) (Hammond, Gur, and Johnson, 2013) and the Ipsen-Mikhailov spectral distance (IMSD) (Jurman et al., 2011), which was designed for comparing biological networks. The GDD is computed as

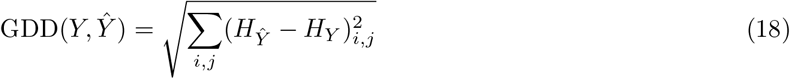

where 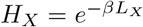 and *L*_*X*_ = *D*_*X*_ −*A*_*X*_ is the graph Laplacian of graph X, and where β is the diffusion parameter, set to 1*/N* for all networks. The IMSD is computed as

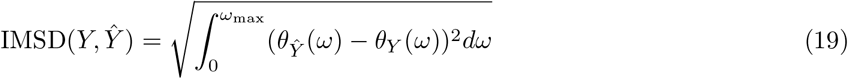

The spectral density *θ* (*·*) of a graph is defined as the sum of Lorentz distributions

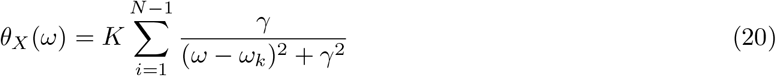

where 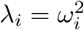 are the non-zero eigenvalues of the Laplacian of *X*, ω_*max*_ is the square root of the largest eigenvalues of *L*_*Y*_, *L*_*ŷ*_,K is the normalization constant, and γ= 0.08 is a scale parameter and is set following the original reference. We use *θ* rather than the original ρ to avoid confusion with the Spearman ρ. For the SHPs, which comprise *N* categorical distributions, we compute the KL divergence

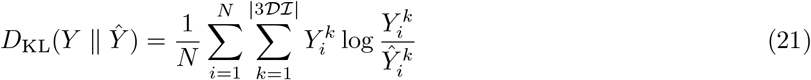

#### Hyperparameters

Classification head weight matrices (*W*_1_, *W*_2_, *W*_3_, *W*_4_, *W*_1*conv*_, *W*_2*conv*_, *W*_3*conv*_, *W*_4*conv*_) all use *d*_*model*_ for the hidden dimension, where for PLANET-MD and PLANET-MD-seq we use *d*_*model*_ = 512, and for PLANET-MD-mini *d*_*model*_ = 128. The sequence embedding dimension from ESM3 is *d*_*seq*_ = 1536, and the structure embedding dimension varies, but the default used in PLANET-MD is *d*_*struct*_ = 1024. For PLANET-MD and PLANET-MD-seq, the transformer encoder contains 8 layers with 8 heads each, while the PLANET-MD-mini transformer encoder has 4 layers with 4 heads. During training, input proteins were cropped to 512 amino acids for GPU memory efficiency. No cropping is done during inference.

We use the default parameters for the Adam optimizer, including *β*_1_ = 0.9, *β*_2_ = 0.99. We tried learning rates 1*e* − 4, 5*e* − 5, 1*e* − 5 and settled on 5 − *e*5 because it yielded the best convergence. Models were trained for 75 epochs and the best model was selected based on validation loss (PLANET-MD, epoch 29; PLANET-MD-seq, epoch 16; PLANET-MD-mini, epoch 39; mdCATH benchmarks, epoch 10) We show training and validation loss curves for each of the three main models in Appendix A.1.

#### Implementation and Runtime

All training, benchmarking, and analysis was performed on the Flatiron Institute internal cluster. All PLANET-MD models were trained on a machine with 96 CPUs, 500GB of memory, and one 1 NVIDIA A100 GPUs with 40GB memory. Training was impelemented using PyTorch v2.5.1 and Lightning v2.5.0 with medium precision. Training the full model on the ATLAS dataset required approximately 5 hours. Training the full model on the mdCATH dataset required approximately 24 hours. Feature extraction using ESM3 embeddings was performed as a pre-processing step and required less than an hour on a single GPU for ATLAS, and *≈*4 hours for mdCATH. Pre-processing of simulation trajectories and computation of derivatives required an additional day for ATLAS and 5 days for mdCATH using only CPU time.

Inference for a protein of 300 residues takes approximately 0.05 seconds on a single A100 GPU, making the model suitable for high-throughput analysis of protein dynamics. For comparison, running a traditional molecular dynamics simulation for a similar-sized protein would require between 24-48 hours on a GPU for a 100ns trajectory, highlighting the significant computational advantage of our approach for estimating protein dynamics.

#### Dyna-1 Calibration

Dyna-1 was originally trained on the mBMRB-Train dataset, which is currently not publicly available. However, they also make available the RelaxDB dataset, which contained NMR backbone relaxation measurements curated for 133 proteins. We randomly sampled six proteins from this dataset–*11080, BLOT5, CHEA, RNASE, CRABP2*, and *SPOOF*–and ran 100ns MD simulations using OpenMM v8.1.2 with the amber14 force field. We chose to run MD only on a small subset of the proteins for calibration due to computational costs of all-atom MD. Initial structures were modeled using Modeller and were solvated in a periodic box with 1 nanometer padding. After energy minimization, NVT equilibriation was run for 200 picoseconds followed by 1 nanosecond of NPT equilibriation at 1 bar and 300K. We used a 2 femtosecond time step and saved the conformation every 10 picoseconds.

These trajectories were used to compute the RMSF for each chain. We then fit a linear regression model using sklearn to predict the simulated RMSF from the Dyna-1 prediction. This yielded the following equation:

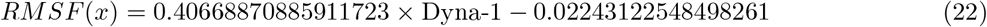

which was used to adjust RMSF predictions for the Dyna-1 model.

### Exponential Scaling of Temperature

We compare PLANET-MD predictions of temperature with a baseline established in Sá Ribeiro and Lima, 2023. They report the following dependency between experimental B-factors (we treat this here as a proxy for RMSF) and temperature:

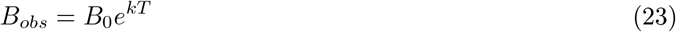

where they fit the parameters *k* = 0.0045*K*^−1^, *B*_0_ = 5 Å. We use this dependency to convert an observed 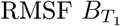 at temperature *T*_1_ = 320*K* to another temperature *T*_2_.

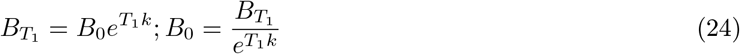

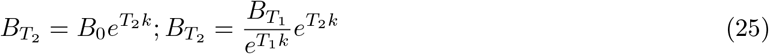

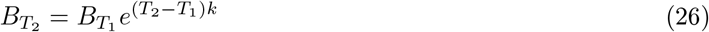

## References

[1] Lalima G. Ahuja, Susan S. Taylor, and Alexandr P. Kornev. “Tuning the “violin” of protein kinases: The role of dynamics-based allostery”. en. In: IUBMB Life 71.6 (2019), pp. 685–696. issn: 1521-6551. doi: 10.1002/iub.2057. url: (visited on 04/09/2025).

[2] Miro A. Astore et al. “Protein dynamics underlying allosteric regulation”. In: Current Opinion in Structural Biology 84 (Feb. 2024), p. 102768. issn: 0959-440X. doi: 10.1016/j.sbi.2023.102768. URL: https://www.sciencedirect.com/science/article/pii/S0959440X23002427 (visited 06/06/2024).

[3] Antoni Beltran et al. “The allosteric landscape of the Src kinase”. In: Science Advances 12.7 (2026), eaea2726.

[4] Tristan Bepler and Bonnie Berger. “Learning the protein language: Evolution, structure, and function”. English. In: Cell Systems 12.6 (June 2021), 654–669.e3. issn: 2405-4712. doi: 10.1016/j.cels.2021.05.017 https://www.cell.com/cell-systems/abstract/S2405-4712(21)00203-9 url: (visited on 08/01/2021).

[5] Mattia Bernetti et al. “Probing allosteric communication with combined molecular dynamics simulations and network analysis”. In: Current Opinion in Structural Biology 86 (2024), p. 102820.

[6] S. Bowerman and J. Wereszczynski. “Detecting Allosteric Networks Using Molecular Dynamics Simulation”. eng. In: Methods in Enzymology 578 (2016), pp. 429–447. issn: 1557-7988. doi:10.1016/bs.mie.2016.05.027.

[7] Stephen K Burley et al. “RCSB Protein Data Bank (RCSB.org): delivery of experimentally-determined PDB structures alongside one million computed structure models of proteins from artificial intelligence/machine learning”. In: Nucleic Acids Research 51.D1 (2023), pp. D488–D508.

[8] Adrienne D Cox et al. “Drugging the undruggable RAS: Mission possible?” In: Nature reviews Drug discovery 13.11 (2014), pp. 828–851.

[9] Peter Eastman et al. “OpenMM 8: molecular dynamics simulation with machine learning potentials”. In: The Journal of Physical Chemistry B 128.1 (2023), pp. 109–116. https://onlinelibrary.wiley.com/

[10] K. Gunasekaran, Buyong Ma, and Ruth Nussinov. “Is allostery an intrinsic property of all dynamic proteins?” en. In: Proteins: Structure, Function, and Bioinformatics 57.3 (Nov. 2004), pp. 433–443. issn: 0887-3585, 1097-0134. doi: 10.1002/prot.20232. url: doi/10.1002/prot.20232 (visited on 11/04/2024).

[11] Gordon G Hammes, Yu-Chu Chang, and Terrence G Oas. “Conformational selection or induced fit: a flux description of reaction mechanism”. In: Proceedings of the National Academy of Sciences 106.33 (2009), pp. 13737–13741.

[12] David K Hammond, Yaniv Gur, and Chris R Johnson. “Graph di”usion distance: A di”erence measure for weighted graphs based on the graph Laplacian exponential kernel”. In: 2013 IEEE global conference on signal and information processing. IEEE. 2013, pp. 419–422.

[13] Matt J Harvey, Giovanni Giupponi, and G De Fabritiis. “ACEMD: accelerating biomolecular dynamics in the microsecond time scale”. In: Journal of chemical theory and computation 5.6 (2009), pp. 1632–1639.

[14] Thomas Hayes et al. Simulating 500 million years of evolution with a language model. en. Pages: 2024.07.01.600583 Section: New Results. July 2024. doi: 10.1101/2024.07.01.600583. url: https://www.biorxiv.org/content/10.1101/2024.07.01.600583v1 (visited on 07/11/2024).

[15] Katherine A Henzler-Wildman et al. “A hierarchy of timescales in protein dynamics is linked to enzyme catalysis”. In: Nature 450.7171 (2007), pp. 913–916.

[16] Jing Huang et al. “CHARMM36m: an improved force field for folded and intrinsically disordered proteins”. In: Nature methods 14.1 (2017), pp. 71–73.

[17] John Jumper et al. “Highly accurate protein structure prediction with AlphaFold”. en. In: Nature 596.7873 (Aug. 2021), pp. 583–589. issn: 0028-0836, 1476-4687. doi: 10.1038/s41586-021-03819-2.https://www.nature.com/articles/s41586-021-03819-2url: (visited on 02/16/2023).

[18] Giuseppe Jurman et al. Biological network comparison via Ipsen-Mikhailov distance. arXiv:1109.0220 [q-bio]. Sept. 2011. doi: 10.48550/arXiv.1109.0220. url: http://arxiv.org/abs/1109.0220 (visited on 05/07/2025).

[19] Dan Kalifa, Eric Horvitz, and Kira Radinsky. “Learning protein representations with conformational dynamics”. In: Bioinformatics 42.5 (2026).

[20] Michel van Kempen et al. “Fast and accurate protein structure search with Foldseek”. en. In: Nature Biotechnology 42.2 (Feb. 2024). Number: 2, pp. 243–246. issn: 1546-1696. doi: 10.1038/s41587-023-01773-0.https://www.nature.com/articles/s41587-023-01773-0url: (visited on 03/01/2024).

[21] Dorothee Kern and Erik RP Zuiderweg. “The role of dynamics in allosteric regulation”. In: Current opinion in structural biology 13.6 (2003), pp. 748–757.

[22] Michael Knudsen and Carsten Wiuf. “The CATH database”. In: Human genomics 4 (2010), pp. 1–6.

[23] Solomon Kullback. Kullback-leibler divergence. 1951.

[24] Cedric Leroux and Georgia Konstantinidou. “Targeted therapies for pancreatic cancer: overview of current treatments and new opportunities for personalized oncology”. In: Cancers 13.4 (2021), p. 799.

[25] Sarah Lewis et al. “Scalable emulation of protein equilibrium ensembles with generative deep learning”. In: Science 389.6761 (July 2025), eadv9817. doi: 10.1126/science.adv9817. url: https://www.science.org/doi/10.1126/science.adv9817 (visited on 08/26/2025).

[26] Xiaotian Liao and Ben Lehner. Allostery is a widespread cause of loss-of-function variant pathogenicity. en. Pages: 2025.06.20.660737 Section: New Results. June 2025. doi: 10.1101/2025.06.20.660737.urlhttps://www.biorxiv.org/content/10.1101/2025.06.20.660737v1 (visited on 06/25/2025).

[27] Zeming Lin et al. “Evolutionary-scale prediction of atomic-level protein structure with a language model”. en. In: Science 379.6637 (Mar. 2023), pp. 1123–1130. issn: 0036-8075, 1095-9203. doi:10.1126/science.ade2574.url:https://www.science.org/doi/10.1126/science.ade2574 (visited on 01/17/2024).

[28] Janet E. Lindsley and Jared Rutter. “Whence cometh the allosterome?” en. In: Proceedings of the National Academy of Sciences 103.28 (July 2006), pp. 10533–10535. issn: 0027-8424, 1091-6490. doi:10.1073/pnas.0604452103.https://pnas.org/doi/full/10.1073/pnas.0604452103 url: (visited on 09/23/2024).

[29] Yikai Liu et al. “ExEnDi”: An Experiment-Guided Di”usion Model for Protein Conformational Ensemble Generation”. In: PRX Life 3.2 (June 2025), p. 023013. doi: 10.1103/PRXLife.3.023013.url:https://link.aps.org/doi/10.1103/PRXLife.3.023013 (visited on 08/27/2025).

[30] Federica Maschietto et al. “MDiGest: A Python package for describing allostery from molecular dynamics simulations”. In: The Journal of Chemical Physics 158.21 (June 2023), p. 215103.issn:0021-9606.doi:10.1063/5.0140453.url:https://doi.org/10.1063/5.0140453(visited on 03/17/2025).

[31] Till Maurer et al. “Small-molecule ligands bind to a distinct pocket in Ras and inhibit SOS-mediated nucleotide exchange activity”. In: Proceedings of the National Academy of Sciences 109.14 (2012), pp. 5299–5304.

[32] Robert T McGibbon et al. “MDTraj: a modern open library for the analysis of molecular dynamics trajectories”. In: Biophysical journal 109.8 (2015), pp. 1528–1532.

[33] Antonio Mirarchi, Toni Giorgino, and Gianni De Fabritiis. mdCATH: A Large-Scale MD Dataset for Data-Driven Computational Biophysics. arXiv:2407.14794 [q-bio]. July 2024. doi: 10.48550/arXiv.2407.14794. url: http://arxiv.org/abs/2407.14794 (visited on 09/19/2024).

[34] Christine A Orengo et al. “CATH–a hierarchic classification of protein domain structures”. In: Structure 5.8 (1997), pp. 1093–1109.

[35] Tatu Pantsar. “The current understanding of KRAS protein structure and dynamics”. In: Computational and structural biotechnology journal 18 (2020), pp. 189–198.

[36] Alexander Rives et al. “Biological structure and function emerge from scaling unsupervised learning to 250 million protein sequences”. In: bioRxiv (Aug. 2019). tex.ids= rives2020scaling, rives2021biological tex.elocation-id: 622803, pp. 622803–622803. doi: 10.1101/622803. url: https://doi.org/10.1101/622803.

[37] Meagan B Ryan and Ryan B Corcoran. “Therapeutic strategies to target RAS-mutant cancers”. In: Nature reviews Clinical oncology 15.11 (2018), pp. 709–720.

[38] Fernando de Sá Ribeiro and Luís Maurício TR Lima. “Linking B-factor and temperature-induced conformational transition”. In: Biophysical Chemistry 298 (2023), p. 107027.

[39] Hugo Schweke et al. “An atlas of protein homo-oligomerization across domains of life”. In: bioRxiv (2023), pp. 2023–06.

[40] Sukrit Singh and Gregory R. Bowman. “Quantifying Allosteric Communication via Both Concerted Structural Changes and Conformational Disorder with CARDS”. In: Journal of Chemical Theory and Computation 13.4 (Apr. 2017), pp. 1509–1517. issn: 1549-9618. doi: 10.1021/acs.jctc.6b01181.url:https://www.sciencedirect.com///doi.org/10.1021/acs.jctc.6b01181 (visited on 04/09/2025).

[41] Samuel Sledzieski and Sonya M. Hanson. “The landscape of machine learning approaches for modeling protein conformational ensembles”. In: Current Opinion in Structural Biology 98 (June 2026), p. 103253. issn: 0959-440X. doi: 10.1016/j.sbi.2026.103253. url: science/article/pii/S0959440X26000357 (visited on 04/29/2026).

[42] Martin Steinegger and Johannes Söding. “Clustering huge protein sequence sets in linear time”. In: Nature communications 9.1 (2018), p. 2542.

[43] Kenichi Suda, Kenji Tomizawa, and Tetsuya Mitsudomi. “Biological and clinical significance of KRAS mutations in lung cancer: an oncogenic driver that contrasts with EGFR mutation”. In: Cancer and Metastasis Reviews 29.1 (2010), pp. 49–60.

[44] Baris E Suzek et al. “UniRef clusters: a comprehensive and scalable alternative for improving sequence similarity searches”. In: Bioinformatics 31.6 (2015), pp. 926–932.

[45] David Van Der Spoel et al. “GROMACS: fast, flexible, and free”. In: Journal of computational chemistry 26.16 (2005), pp. 1701–1718.

[46] Mihaly Varadi et al. “AlphaFold Protein Structure Database: massively expanding the structural coverage of protein-sequence space with high-accuracy models”. In: Nucleic Acids Research 50.D1 (Jan. 2022), pp. D439–D444. issn: 0305-1048. doi: 10.1093/nar/gkab1061. url: https://doi.org/10.1093/nar/gkab1061 (visited on 05/09/2025).

[47] Hannah K. Wayment-Steele et al. Learning millisecond protein dynamics from what is missing in NMR spectra. en. Pages: 2025.03.19.642801 Section: New Results. Mar. 2025. doi: 10.1101/2025.03.19.642801. url: https://www.biorxiv.org/content/10.1101/2025.03.19.642801v1 (visited on 03/20/2025).

[48] Hannah K. Wayment-Steele et al. “Predicting multiple conformations via sequence clustering and AlphaFold2”. en. In: Nature 625.7996 (Jan. 2024). Number: 7996, pp. 832–839. issn: 1476-4687. doi: 10.1038/s41586-023-06832-9. url: https://www.nature.com/articles/s41586-023-06832-9 (visited on 02/01/2024).

[49] Chenchun Weng et al. “The energetic and allosteric landscape for KRAS inhibition”. en. In: Nature 626.7999 (Feb. 2024), pp. 643–652. issn: 1476-4687. doi:10.1038/s41586-023-06954-0. url:https://www.nature.com/articles/s41586-023-06954-0(visited on 03/13/2024).

[50] Brian M Wolpin et al. “Daraxonrasib in previously treated advanced RAS-mutated pancreatic cancer”. In: New England Journal of Medicine 394.18 (2026), pp. 1790–1802.

[51] Ruidong Wu et al. “High-resolution de novo structure prediction from primary sequence”. In: BioRxiv(2022), pp. 2022–07.

[52] Mai Yamauchi et al. “Assessment of colorectal cancer molecular features along bowel subsites challenges the conception of distinct dichotomy of proximal versus distal colorectum”. In: Gut 61.6 (2012), pp. 847–854.

