## Appendix for "PLANET-MD: Ultra-fast Proteome-scale Prediction of Allosteric Networks in Proteins"

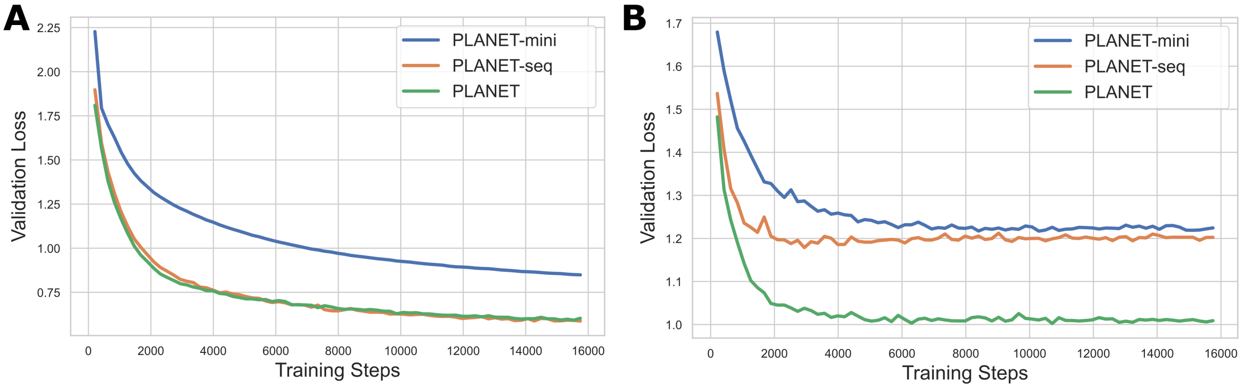


Figure A1: **PLANET-MD Train (A) and Validation (B) Loss Curves**

### A Appendix

#### A.1 Training Curves

In Figure A1, we show training and validation loss curves for the three variants of PLANET-MD. All models are converged, but the full (structure inclusive) model converges at a lower loss than the two sequence-only models.

#### A.2 Performance on MDCath Dataset

Data preparation for mdCATH followed that of ATLAS. We likewise generated training, validation, and testing splits using FoldSeek to ensure diverse structures between sets. As mdCATH contains simulations at five different temperatures, both replicates and different temperatures of the same chain were placed in the same set. The training/validation/test sets then contained 4053/509/622 chains, with 25 samples of each (five temperatures times five replicates, Figure 5a). Simulation temperature was provided as an input to the model during training. Baseline comparisons with Dyna-1 and BioEmu were at the lowest temperature (320K) which is closest to the native biological temperature. In Table A1, we show the performance of each method. As with ATLAS, PLANET-MD outperforms BioEmu and Dyna-1 on all metrics except IMSD.

#### A.3 KRAS Case Study

In a case study, we applied PLANET-MD to model dynamics in *KRAS*, including building an allosteric network, identifying clusters of correlated motion, and performing an *in silico* deep mutational scan to identify residue with a high folding $\Delta\Delta G$. We downloaded the AlphaFold structure of *KRAS* (UniProt: P01116)

Table A1: **Performance on mdCATH Benchmarks.** We compare leading methods for predicting residue flexibility (RMSF), correlation (GCC-LMI), and structure heterogeneity (SHP) on mdCATH. Lower is better for all methods except Spearman $\rho$. The best performance for each is **bolded.** Ground truth for all is at the lowest temperature of 320K.

| Model | RMSF | | GCC-LMI | | SHP | Model Information | | |
|---|---|---|---|---|---|---|---|---|
| | **RMSE** | **Spear.** $\rho$ (↑) | **GDD** | **IMSD** | **KL-Div** | **Seq.** | **Struct.** | **Params.** |
| PLANET-MD | **0.113** | **0.796** | **0.614** | 2.329 | **0.865** | ✓ | ✓ | 30.2M |
| BioEmu (100 samples) | 0.157 | 0.760 | 0.739 | **1.929** | 1.684 | ✓ | ✗ | 31.2M |
| Dyna-1 (calibrated) | 0.218 | 0.351 | - | - | - | ✓ | ✓ | 188.9M |

from AlphaFoldDB and used this as our starting structure for simulation, as well as the input structure to PLANET-MD. We performed a 100ns simulation of *KRAS* to be our "ground truth" dynamics.

Predictions were made using the full PLANET-MD model, trained on ATLAS. Using the predicted $C\alpha$ distances, we masked the predicted $GCC - LMI$ at 6 angstroms and constructed an allosteric network. We clustered this network using the Girvan-Newman algorithm as implemented in the networkx package with $k = 5$ to identify communities within the allosteric network. In Figure A3A, we show the AlphaFold structure of *KRAS* colored with each of the clusters. We perform an in-house simulation of *KRAS* to treat as a ground truth using OpenMM v8.1.2 with the amber14 force field. We used the structure of KRAS downloaded from AlphaFoldDB (P01116). The initial structures was modeled using Modeller and was solvated in a periodic box with 1 nanometer padding. After energy minimization, we ran 200 picoseconds of NVT equilibriation followed by 1 nanosecond of NPT equilibriation at 1 bar and 300K. We ran 100ns of simulation using a 2 femtosecond time step, and saved the conformation every 10 picoseconds. In Figure A3B, C, and D, we show the true dynamic properties (simulation derived) for *KRAS*, compared to the PLANET-MD predictions. We find a strong agreement between predicted and simulated dynamics. Each metric is also colored by cluster of the allosteric network; the predicted RMSF correctly identifies the peak in the binding pocket (blue cluster) and the increasing RMSF in the C-terminal tail (yellow cluster).

We perform an *in silico* deep mutational scan of *KRAS* and compare this to the experimental DMS from Weng et al. (Weng et al., 2024). We mutate each position of *KRAS* to all other potential amino acids. For each mutant, we compute the importance of the residue for dynamic information flow in the network by masking the predicted $GCC - LMI$ at 8 angstroms using the predicted $C\alpha$ distances, then computing the betweenness centrality of the mutated residue in the allosteric network. We used the betweenness centrality as implemented in the networkx package v3.4.2. Here, we are using the full PLANET-MD model, which requires a structure input. For the sake of computational efficiency we chose to use to use the wild type

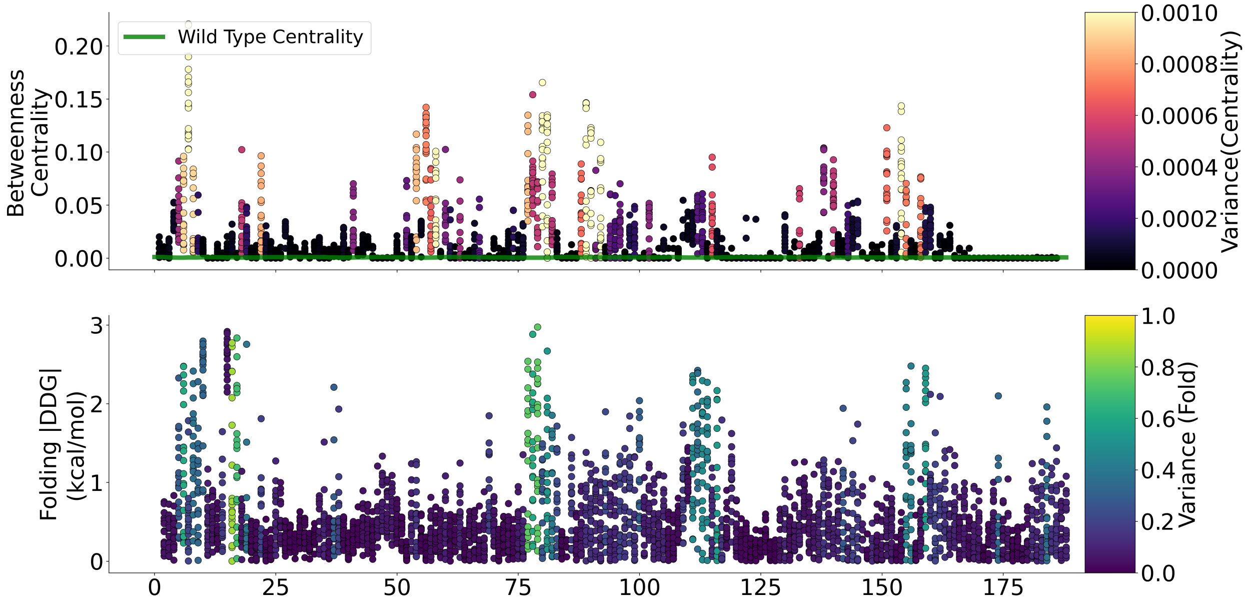


Figure A2: ***KRAS* DMS with PLANET-MD-Seq.**

structure with each mutant sequence; a potentially more accurate approach would be to re-fold each mutant sequence with AlphaFold, although this incurs a significantly higher runtime cost. In Figure A2, we show the results of a DMS using PLANET-MD-seq.

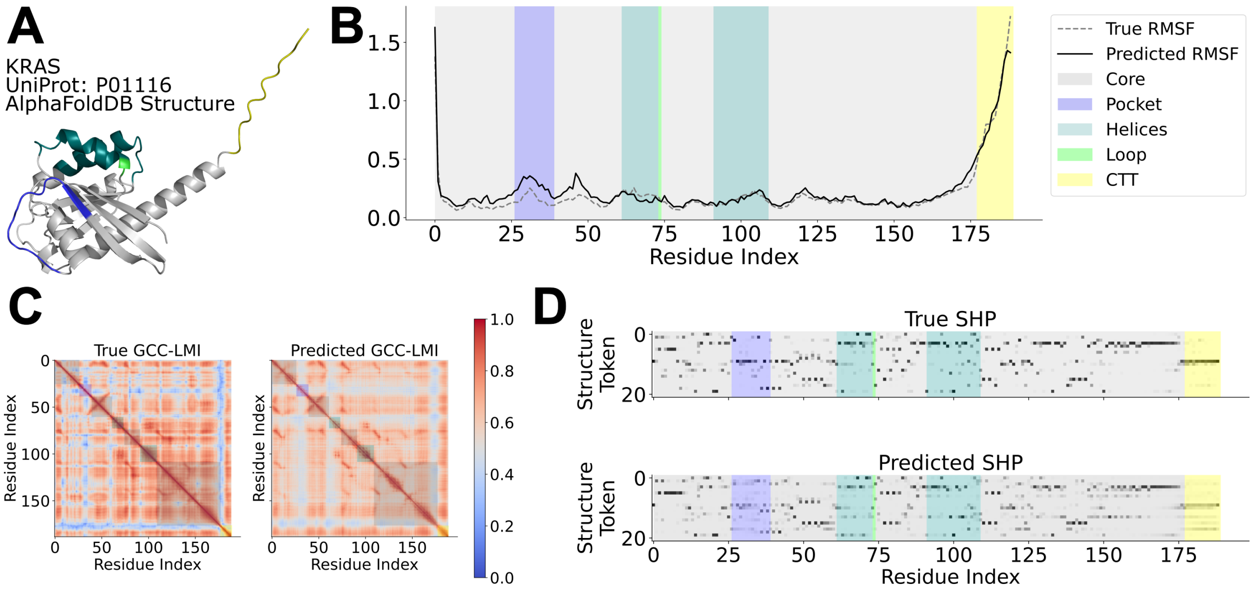


Figure A3: **Sample PLANET-MD predictions for *KRAS*.** **(A)** AlphaFoldDB structure for *KRAS*, colored by Girvan-Newman cluster. **(B)** True (dashed, gray) vs. predicted (solid, black) RMSF. **(C)** True (left) vs. predicted (right) GCC-LMI. **(D)** True (top) vs. predicted (bottom) SHP.
